# Prolactin-induced AMPK stabilizes alveologenesis and lactogenesis through regulation of STAT5 signaling

**DOI:** 10.1101/2022.02.15.480514

**Authors:** Shyam Lal Jinagal, Pragati Shekhar, Kailash Chandra, Srinivas Abhishek Mutnuru, Narendrakumar Ramanan, Marc Foretz, Benoit Viollet, Ramray Bhat, Annapoorni Rangarajan

## Abstract

AMP-activated protein kinase (AMPK) is an evolutionarily conserved serine/threonine kinase that regulates energy homeostasis at cellular and organismal levels. It has been shown to affect several steps of breast cancer progression in a context-dependent manner. However, its role in normal mammary gland development and physiology remains ill-explored. Here, we show that AMPK expression and activity increased within murine mammary epithelia from puberty to pregnancy with highest levels during lactation, and then declined during involution. In *ex vivo* cultures of mammary epithelial cells (MECs) in organotypic scaffolds, treatment with lactogenic hormone prolactin (PRL) enhanced AMPK expression and activity. To understand the role of AMPK on mammary morphogenesis *in vivo*, we generated mice with conditional knockout of AMPKα isoforms α1 and α2 (AMPKα KO) in MECs. AMPKα KO mammary glands showed accelerated alveolar development with increased epithelial content of both luminal and myoepithelial lineages, suggestive of hyperproliferation. AMPKα KO mice also showed elevated beta-casein expression during pregnancy and lactation. These observations were phenocopied upon treatment of *ex vivo* cultivated wild-type MECs with a cognate AMPK inhibitor. AMPKα null MECs showed increased phosphorylated STAT5 which is known to drive alveologenesis downstream of prolactin signaling. Our study identifies a novel interplay between AMPK and PRL-STAT5 signaling that determines mammary alveologenesis and differentiation.

## Introduction

The development of the mammary gland begins embryonically, but its morphogenesis is predominantly completed postnatally in response to various hormonal cues (Richert et al., 2000; Sternlicht, 2005). Under the influence of estrogen and progesterone, the ductal epithelial cells proliferate, invade, and form secondary and tertiary branches, filling the mammary fad pad. The development of mammary alveoli during pregnancy and lactation requires epithelial cell proliferation and coordinated differentiation with maintenance of tissue and cell polarity. During lactation, a functional lobuloalveolar unit consisting of milk-producing inner luminal cells and outer layer of myoepithelial cells is formed (Horseman, 1999; Tucker, 1979). The mammary gland requires abundant energy during lactation to maintain basal metabolism and synthesize milk for neonates (Sampson and Jansen, 1984). Such energy demands are essential to regulate cell survival and function during mammary gland development (Wu et al., 2020). Prolactin (PRL) is a lactogenic hormone that plays a key role in mediating mammary alveolar growth and differentiation (Ormandy et al., 1997). PRL binds to its receptor (PRLR) and triggers its dimerization and phosphorylation by receptor-associated tyrosine kinase Janus kinase 2 (JAK2). JAK2 further recruits SH2 domain-containing signal transducers and activators of transcription (STAT5a and STAT5b) proteins to the receptor complex and phosphorylates them. Activated STAT5a and STAT5b translocate into the nucleus as a dimer, bind to GAS sequences (TTCNNNGAA), and induce transcription of target genes that promote proliferation, differentiation, and lactogenesis (Cui et al., 2004; Teglund et al., 1998; Wagner et al., 2004).

The AMP-activated protein kinase (AMPK) is a sensor of cellular energy status that gets activated in response to stresses that lead to low ATP: AMP and ATP: ADP ratios (Carling, 2019; Hardie and Hawley, 2001; Hardie et al., 1999; Kahn et al., 2005). AMPK is a heterotrimeric Ser/Thr kinase consisting of one catalytic subunit α and two regulatory subunits β and γ. While α and β subunits have two isoforms (α1, α2) and (β1, β2) respectively, the γ subunit has three isoforms (γ1, γ2, and γ3). Several physiological stresses such as ischemia, hypoxia, nutrient deprivation, and matrix-detachment activate AMPK (Fung et al., 2008; Hindupur et al., 2014; Marsin et al., 2000, 2002; Minokoshi et al., 2004; Ng et al., 2012). It plays a central role in bringing about energy homeostasis by inhibiting anabolic processes that consume ATP while turning on catabolic processes (Hardie et al., 2012). In addition to its central role in metabolism, AMPK regulates diverse cellular processes such as cell shape and polarity, cell proliferation and survival, autophagy, and gene transcription through crosstalk with several signaling pathways such as the Wnt/B catenin, GSK3 beta, PEA15, mTOR, and YAP/TAZ (Mihaylova and Shaw, 2011). Some cellular effects of AMPK have also been demonstrated in mammary epithelial cells (MECs) (Guo et al., 2020; Li et al., 2020; Wang et al., 2019). However, how such effects manifest at the tissue scale to modulate the morphology and physiology of the mammary gland is not well understood. *In vitro* studies using bovine MECs have shown an association between AMPK and JAK2-STAT5 pathway under glucose-deprived conditions (Wu et al., 2020; Zhang et al., 2018), although such associations have not yet been confirmed *in vivo*.

In this study, we hypothesized that AMPK maintains cellular energy homeostasis during mammary gland development and regulates epithelial growth and differentiation. Since wholebody germline AMPKα double knockout (α1^-/-^, α2^-/-^) mice are embryonically lethal at ~E10.5 days (Viollet et al., 2009), we generated conditional knockout mice in which AMPKα1 and AMPKα2 were deleted in MECs via an MMTV promoter-driven Cre transgenic mouse line, in which Cre recombinase expression is seen in MECs from puberty onwards (Wagner et al., 1997). This study revealed an overall increase in epithelial content and milk synthesis in AMPK null mammary glands, along with an increase in STAT5 phosphorylation and its corresponding target genes. Our study establishes a novel role for AMPK signaling in regulating alveologenesis during lactation through a transductional crosstalk with PRL-JAK2-STAT5 pathway, thereby, integrating metabolic signaling network with hormonal pathways.

## Results

### AMPK expression and activity are highest in lactating mammary gland epithelia

To investigate the status of AMPK signaling during mammary gland development, we first analysed the steady-state expression of AMPK *in vivo*. We harvested murine inguinal mammary glands at five different developmental stages (puberty week 5, adult week 10, midpregnancy day 13-15, lactation day 7, and involution day 7) (Figure S1A) and performed immunoblotting using an AMPKα-specific antibody that recognizes both AMPKα1 and α2 isoforms. An increase in AMPKα levels was observed from mid-pregnancy, peaking at lactation and declining at involution (Figure 1A). Since phosphorylation of AMPKα at Thr 172 by upstream kinases leads to its full activation (Hawley et al., 1996), we determined the functional status of AMPK signaling by measuring pAMPKα^T172^. Similar to AMPKα expression, pAMPKα^T172^ levels increased at pregnancy and peaked during lactation (Figure 1A). Since AMPK α1 and α2 isoforms are expressed in a tissue-specific manner (Quentin et al., 2011; Viollet et al., 2009), we investigated their expression status in murine mammary glands using AMPKα1 and AMPKα2 isoform-specific antibodies. Immunoblotting revealed expression of both the isoforms within lactating mammary glands (Figure 1B). These expression and activity patterns of AMPK was further corroborated by immunohistochemical (IHC) staining that revealed highest levels of AMPKα and pAMPKα^T172^ within lactating alveolar MECs compared with epithelia at other stages (Figure 1Ci-x) (indicated by arrowheads in Figure 1Ciii, 1Civ, and 1Cviii-1Cx) and with sparse staining of pAMPKα^T172^ in the surrounding adipocytes during pregnancy and involution stages (indicated by asterisk in Figure 1Cviii and 1Cx). Since activated AMPK phosphorylates its bona fide substrate Acetyl-CoA Carboxylase (ACC) at Ser79 (Winder and Hardie, 1996), we additionally performed immunohistochemistry for pACC^Ser79^. Similar to pAMPKα^T172^ expression, pACC^Ser79^ staining revealed its highest expression during pregnancy and lactation (Figure 1Cxi-1Cxv). Based on the expression analyses, our observations suggested a potential morphogenetic role for AMPK during pregnancy and lactation.

**Figure 1.**
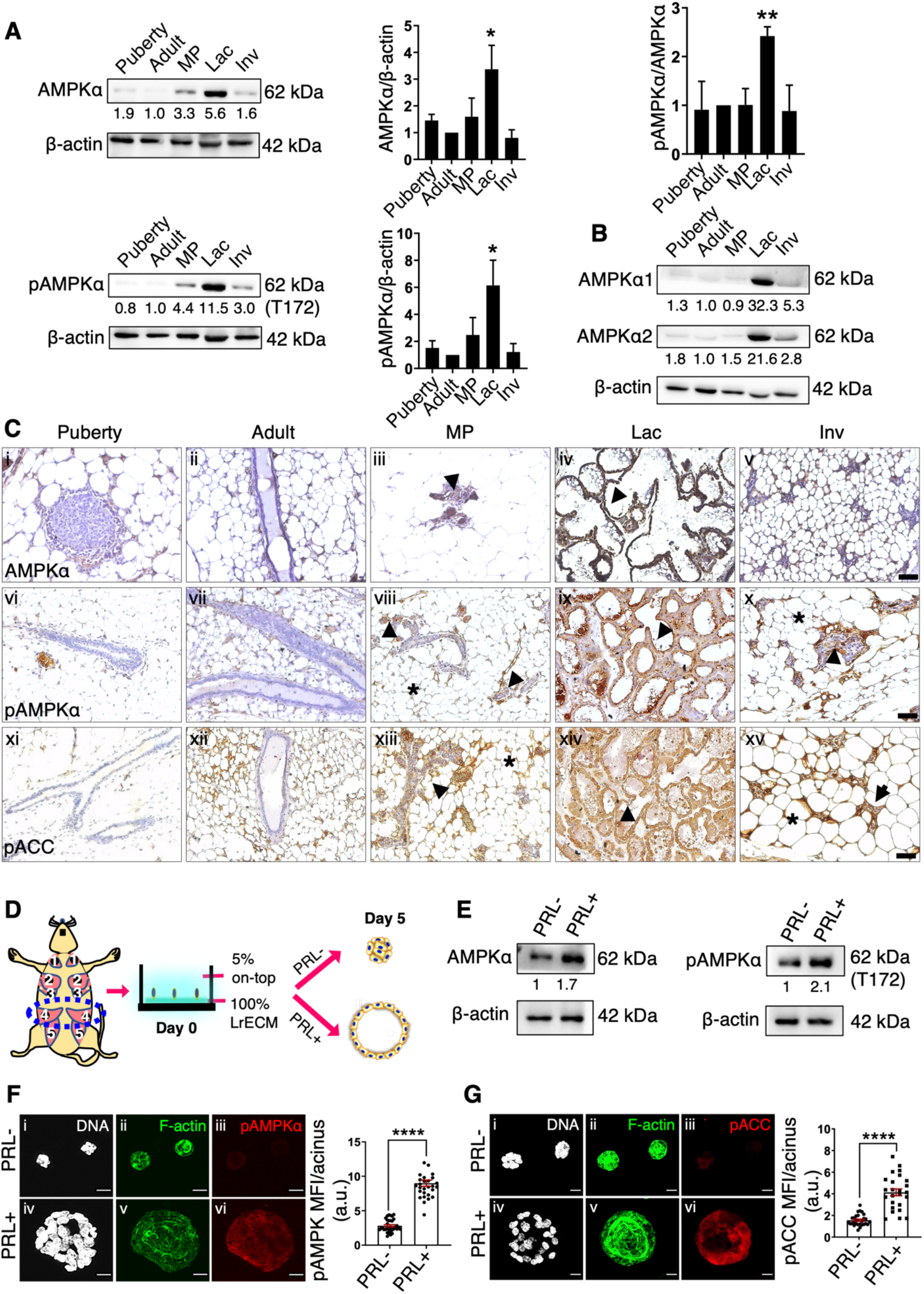
Expression and activity of AMPK across different stages of mammary gland development. (A and B) Representative immunoblots for specified proteins at various stages of mammary gland development in C57/BL6J mice: puberty (5 weeks), adult (10 weeks), MP (mid-pregnant), Lac (day 7 lactation), and Inv (day 7 involution). AMPKα and pAMPKα^T172^ were probed using the same lysate. Graphs represent densitometric quantification of immunoblots; error bars, mean ± SEM; *n* = 4. \**P*<0.05, \*\**P*<0.01, unpaired Student’s *t*-test. (C) Micrographs of mammary gland sections stained for AMPKα (i-v), pAMPKα^T172^ (vi-x), and pACC^Ser79^ (x-xv) (brown) and DNA with hematoxylin (blue). Arrowhead indicates stained epithelial cells, and asterisk indicates stained stromal cells; scale bar: 50 *μ*m. (D) Schematic depiction of *ex vivo* culture of primary mammary epithelial cells (MECs) on-top of LrECM scaffolds in the presence (+) and absence (-) of prolactin (PRL). (E-G) Representative immunoblots (E) and laser confocal micrographs of immunostaining (F and G) of MECs cultured on-top of LrECM as described in the schematic in D; *n* = 3. Micrographs show maximum intensity projection (MIP) of immunofluorescence signals from pAMPKα^T172^ or pACC^Ser79^ (red), staining of F-actin with phalloidin (green), and DNA with Hoechst (white); scale bar: 10 *μ*m. Representative scatter plots with bar graphs show the quantification of immunofluorescence signals (MIP) from pAMPKα^T172^ (F) and pACC^Ser79^ (G) per acinus normalized to F-actin; error bars, mean ± SEM; *n* = 3. \*\*\*\**P*<0.0001. (See also supplementary figure 1).

Prolactin (PRL) signaling through prolactin receptor (PRLR) majorly regulates mammary lobuloalveolar growth and lactogenesis during pregnancy and lactation (Ormandy et al., 1997). Since we observed an induction of AMPK expression and activity at lactation, we asked whether alveologenesis involves signaling crosstalk between PRL and AMPK. To address this, we resorted to *ex vivo* organotypic model of MEC culture in which exogenously added PRL stimulates alveologenesis resulting in acini formation (Gouilleux et al., 1994; Watson and Burdon, 1996), and investigated the status of AMPK. We cultured MECs on laminin-rich ECM (LrECM) scaffold for five days and observed multicellular clusters with lumen in the presence of PRL (Figure 1D). Further, the PRL-treated clusters had increased acinar area and cell number relative to vehicle-treated controls, as well as expressed beta-casein – a marker of lactogenic differentiation (Figure S1B-S1D). These observed effects of PRL on lumenogenesis within acinar-like structures are consistent with earlier reports (Barcellos-Hoff et al., 1989; Streuli et al., 1991). We next investigated the status of AMPK in response to prolactin induction. Immunoblot analyses of such PRL-treated clusters showed a significant increase for AMPKα and pAMPKα^T172^ upon PRL treatment (Figure 1E and S1E). Levels of pAMPKα^T172^ and pACC^Ser79^ and their localization within mammary epithelia were additionally validated using fluorescent immunocytochemistry. Maximum intensity projections (MIPs) revealed expression in all cells within the acini, and quantification of fluorescence signals showed a significant increase in pAMPKα^T172^ and pACC^Ser79^ staining per acinus in the presence of PRL compared with vehicle-treated control acini (Figure 1F and 1G). To the contrary, PRL failed to induce AMPK expression and activity in MECs cultured as monolayers (Figure S1F and S1G). This is in keeping with the requirement of a suitable 3-dimensional environment for proper signaling through PRLR (Xu et al., 2009). These results further suggest that the upregulation of AMPK expression and signaling during pregnancy and lactation is likely driven by PRL signaling.

### Generation of AMPKα conditional knockout mouse model

To better understand the role of AMPK in mammary gland development, we generated an AMPK conditional knockout mouse model using the Cre-LoxP system (Sauer and Henderson, 1988). The *PRKAA1* and *PRKAA2* genes encode for the catalytic subunit of AMPK, AMPKα isoforms α1 and α2, respectively. Since we observed the expression of both isoforms in murine mammary epithelia (Figure 1B), we conditionally deleted both *PRKAA1* and *PRKAA2* genes in the mammary gland epithelium by crossing MMTV-Cre transgenic mice, which express Cre in mammary epithelial cells (Wagner et al., 1997, 2001), with mice carrying floxed *PRKAA1* and *PRKAA2* alleles (AMPKα1^fl/fl^, AMPKα2^fl/fl^) (Figure 2A). AMPK heterozygous knockout animals (AMPKα1^+/-^, AMPKα2^+/-^) produced in the F1 generation were further back-crossed with AMPK floxed (AMPKα1^fl/fl^, AMPKα2^fl/fl^) mice for generation of homozygous double-knockout mice (AMPKα1^-/-^, AMPKα2^-/-^) (Figure S2A), hereafter referred to as AMPK KO mice. To detect *PRKAA1* and *PRKAA2* alleles with and without LoxP and the presence of *Cre* allele, we performed PCR genotyping using specific primers (see Methods, Table 1). In the AMPK KO mice, we observed *PRKAA1* floxed allele and *PRKAA2* floxed allele at 682 bp and 250 bp, respectively, in mouse tail DNA and knockout *PRKAA1* and *PRKAA2* alleles at 348 bp and 600 bp, respectively, in mammary gland genomic DNA, along with the *Cre* transgene band at 400 bp position in agarose gel (Figure 2B). Immunoblot analysis of protein lysate obtained from day 1 lactating mammary gland tissue from AMPK KO mice showed the presence of Cre recombinase protein (Figure 2C). Immunoblotting on mammary glands obtained from control and AMPK KO lactating mice (Figure 2D), as well as in MECs derived from mid-pregnant AMPK KO mice (Figure S2B), showed a significant reduction in AMPKα levels. As observed earlier (Zarrinpashneh et al., 2006), a band corresponding to a truncated AMPKα2 (as a result of deletion of catalytic domain encoded by nucleotides corresponding to 189-260 amino acids) could be seen on the blots from the AMPK KO mammary gland lysates. Immunofluorescence staining of AMPKα of day 1 lactating mammary gland from AMPKα KO mice showed an absence of detectable AMPKα in the mammary epithelia of the AMPKα KO mice (Figure 2E). Immunostaining also revealed that the deletion of AMPKα impaired the phosphorylation of its downstream substrate ACC at Ser79 in mammary epithelia (Figure 2F). Pups were born at the expected Mendelian ratio and were indistinguishable from control mice at birth in terms of weight and morphology.

**Figure 2.**
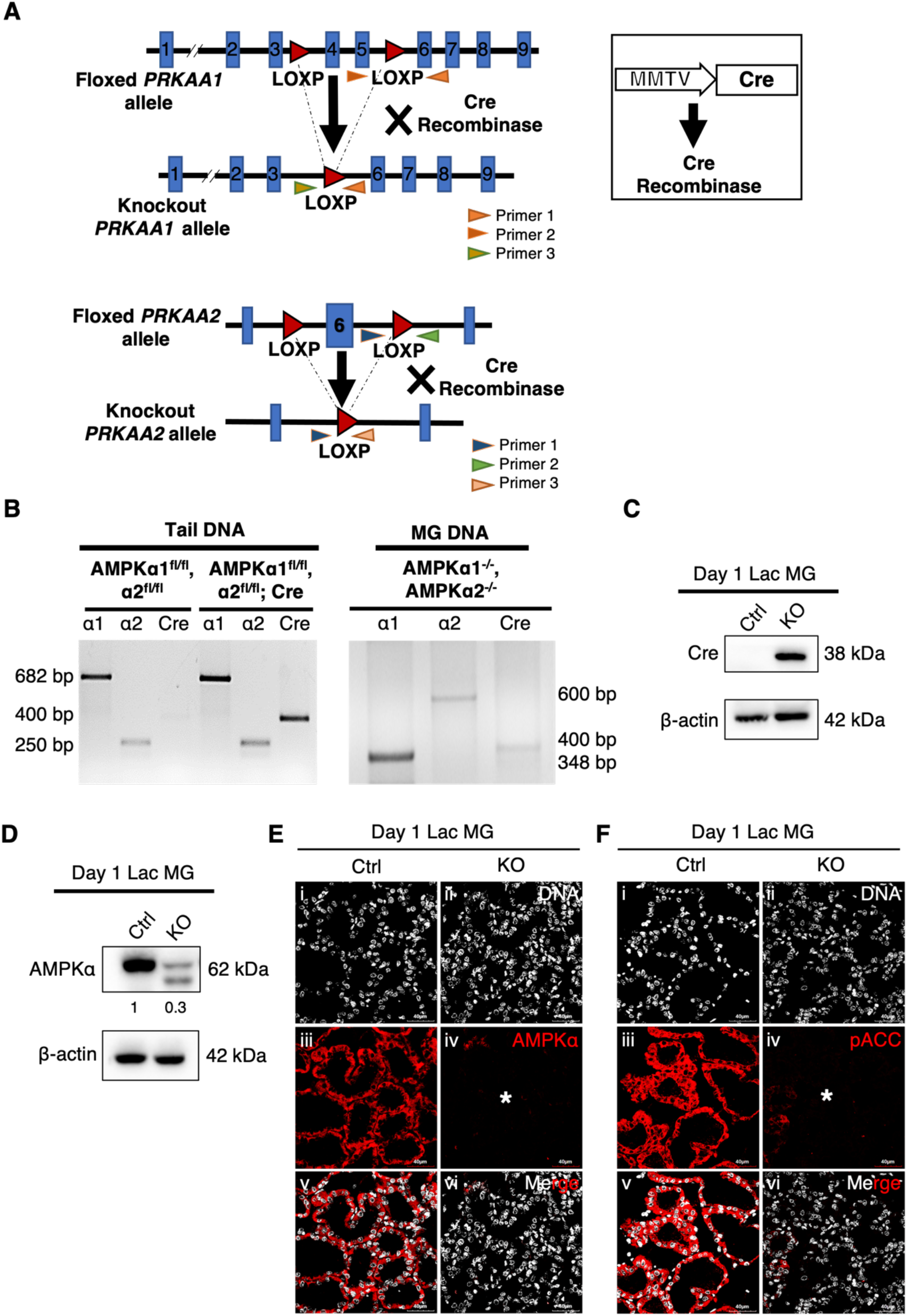
Generation of AMPKα conditional knockout mice. (A) Schematic representation of the strategy used for conditional deletion of *PRKAA1* (AMPKα1) and *PRKAA2* (AMPKα2) in mammary gland epithelium. (B) PCR analysis (using primer sets detailed in methods and indicated in the schematic) shows *PRKAA1* floxed allele and *PRKAA2* floxed allele at 682 bp and 250 bp, respectively, in mouse tail DNA and knockout *PRKAA1* and *PRKAA2* alleles at 348 bp and 600 bp, respectively, in mammary gland (MG) genomic DNA, along with the *Cre* transgene band at 400 bp. (C and D) Representative immunoblots for specified proteins in control and AMPK KO mammary glands at day 1 of lactation. (E and F) Laser confocal micrographs of control and AMPK KO mammary gland sections probed for specified proteins at day 1 of lactation show maximum intensity projection (MIP) of immunofluorescence signals from AMPKα (E) and pACC^Ser79^ (F) (red) and staining of DNA with Hoechst (white). Asterisk indicates AMPK KO mammary epithelium; scale bar: 40 μm. (See also supplementary figure 2).

**Table 1:**
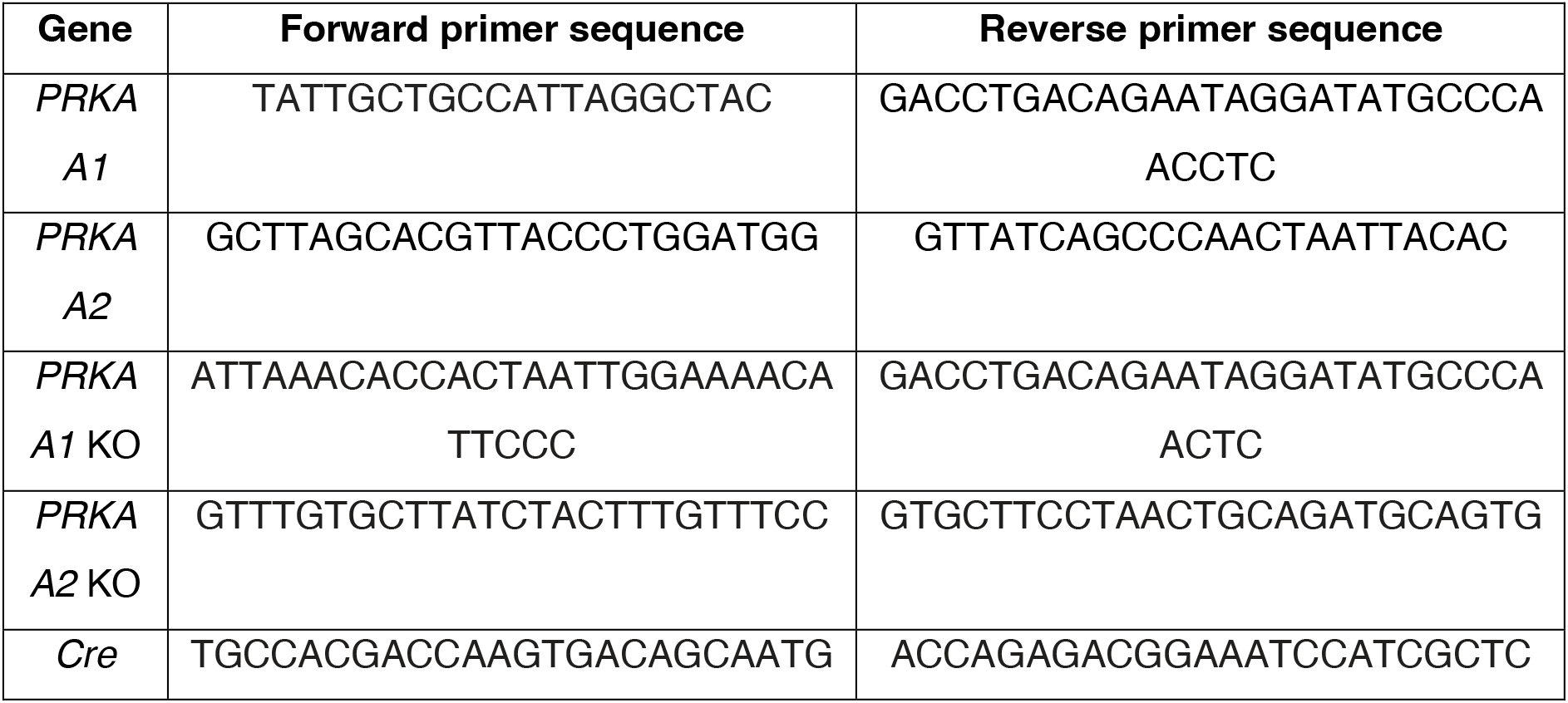
List of primers used in the study

### Deletion of AMPK results in morphological and histological alterations during pregnancy and lactation

We next evaluated the effect of AMPK deletion on mammary gland morphogenesis during pregnancy and lactation, wherein we had observed maximum expression and activity of AMPK (Figure 1A-1C). On day 13.5 of pregnancy, carmine-stained AMPK KO mammary gland whole mounts were observed to be larger in size and showed greater parenchymal density than wildtype controls, as shown and highlighted by asterisk in Figure 3Ai-Aiv. Hematoxylin and Eosin (H&E) staining confirmed increased epithelial content and more lobuloalveolar-like structures in AMPK KO gland sections compared to controls (Figure 3Av and 3Avi). Immunoblot analysis with lysates of day 13.5 pregnant mammary glands for CK18 (a marker for luminal epithelia) and CK14 (a marker for myoepithelia) revealed increased levels for both upon depletion of AMPK (Figure 3B and 3C), suggesting both lineages contribute to the increased parenchymal content when AMPK is knocked out. Similar to the effect of AMPK knockout on mammary gland morphology in pregnancy, glands at day 1 of lactation showed increased epithelial content (visualized through carmine and H&E staining) (Figure 3Di-3Dvi). Quantitative analysis showed a significant increase in epithelial content and reduced stromal content (Figure 3E and 3F). Increased epithelial proportion was further confirmed through increased levels of CK18 and CK14, using immunoblotting of AMPK KO lactating mammary glands (Figure 3G, S3A, 3H, and S3B). Such morphogenetic changes were also observed on day 7 of lactation in AMPK null lactating mammary glands (Figure S3C). Interestingly, day 14 lactating mammary glands did not show any significant changes in epithelial and stromal area; however, an increased number of cells per field was maintained in AMPK deficiency (Figure S3D). This suggests that mammary fat-pad is completely filled at day 14 of lactation in both wild-type and AMPK KO conditions, and AMPK deletion does not cause further changes in the overall epithelial content.

**Figure 3.**
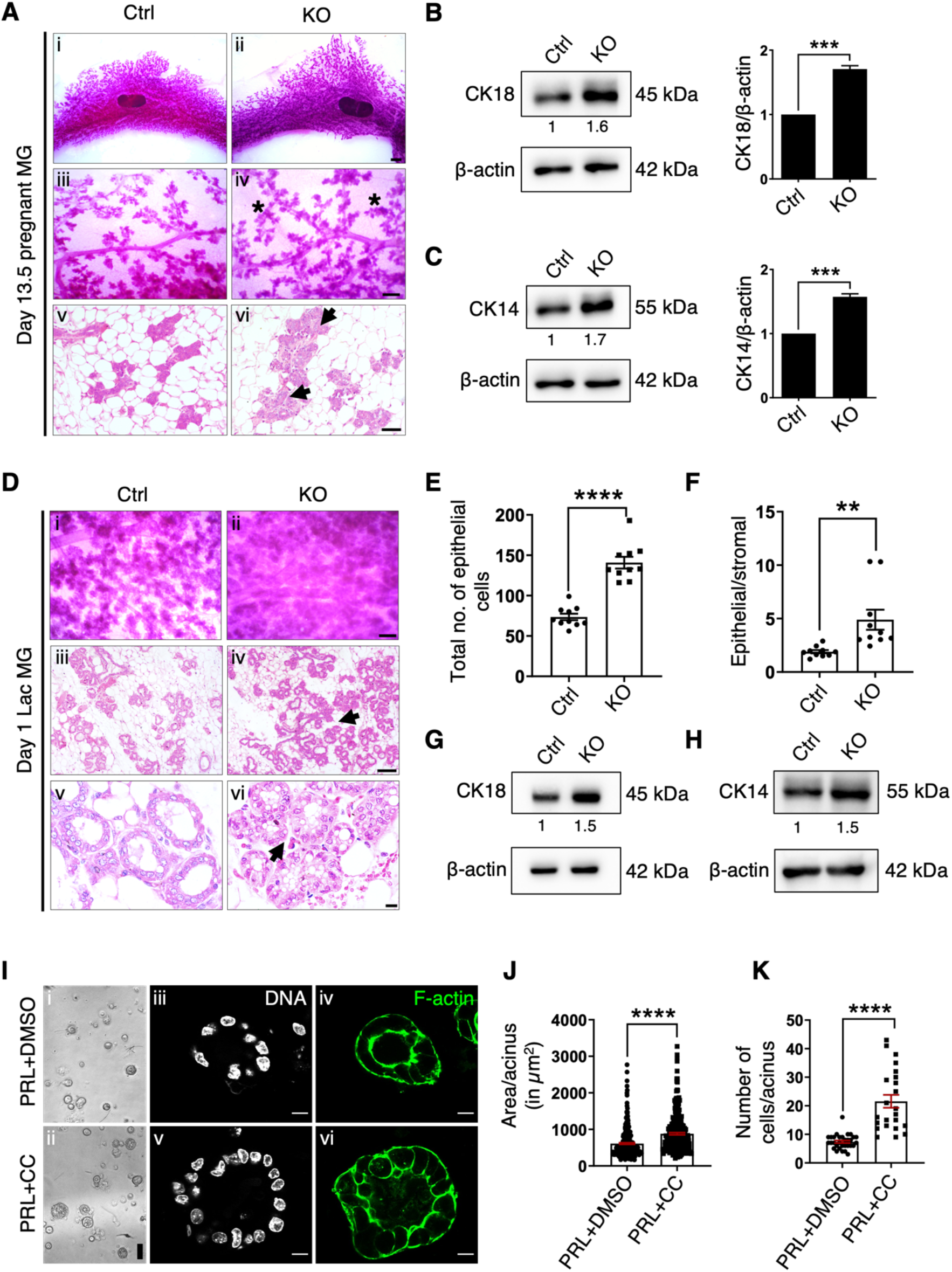
Effect of AMPK ablation on lobuloalveolar morphogenesis. (A) Micrographs of carmine-stained whole mounts (i-iv) and hematoxylin and eosin (H&E) stained sections (v and vi) of control and AMPK KO mammary glands at day 13.5 of pregnancy. Asterisk indicates bigger lobuloalveolar structures (iv) and arrow indicates increased epithelial content (vi) in AMPK KO mammary glands; scale bars: 1 mm (i and ii), 250 *μ*m (iii and iv) and 50 *μ*m (v and vi). (B and C) Representative immunoblots for specified proteins in control and AMPK KO mammary glands at day 13.5 of pregnancy. Graphs represent densitometric quantification of immunoblots; error bars, mean ± SEM; *n* = 3. \*\*\**P*<0.001, unpaired Student’s *t*-test. (D) Micrographs of carmine-stained whole mounts (i and ii) and hematoxylin and eosin (H&E) stained sections (iii-vi) of control and AMPK KO mammary glands at day 1 of lactation. Arrow indicates increased epithelial content (iv and vi) in AMPK KO mammary glands; scale bars: 250 *μ*m (i and ii), 50 *μ*m (iii and iv), and 20 *μ*m (v and vi). (E and F) Representative scatter plots with bar graphs show the quantification of total number of epithelial cells per field and epithelial/ stromal content in control and AMPK KO mammary glands at day 1 of lactation (at least 10 random fields per gland); error bars, mean ± SEM; *n* = 3. \*\**P*<0.01, \*\*\*\**P*<0.0001, unpaired Student’s *t*-test. (G and H) Representative immunoblots for specified proteins in control and AMPK KO mammary glands at day 1 of lactation. (I) Micrographs of phase-contrast (i and ii) and laser confocal middle z-stack (iii-vi) show MECs cultured on-top of LrECM for 5 days in the presence of PRL treated with vehicle control DMSO (i, iii and iv) and compound C (CC) (ii, v and vi), staining of F-actin with phalloidin (green), and DNA with Hoechst (white); scale bar: 50 μm (i and ii) and 10 μm (iii-vi). (J and K) Representative scatter plots with bar graphs show the quantification of area and total number of cells per acinus; error bars, mean ± SEM; *n* = 3. \*\*\*\**P*<0.0001, unpaired Student’s *t*-test. (See also supplementary figure 3).

Furthermore, to investigate the functional role of AMPK in lactating MECs, we studied the effect of AMPK inhibition on acinar morphogenesis using compound C (abbreviated as CC), a pharmacological inhibitor of AMPK (Zhou et al., 2001). The inhibition of AMPK by CC was confirmed by observing a decrease in the levels of pAMPKα^T172^ using immunoblotting (Figure S3E). Whereas control MECs cultured in the presence of PRL and DMSO (vehicle for CC) underwent normal morphological differentiation and formed well-organized acini (Figure 3Ii, 3Iiii, and 3Iiv), those treated with CC led to the formation of acini that were comparatively larger in size and displayed impaired morphology (Figure 3Iii, 3Iv, and 3Ivi). Quantitative analysis revealed that the area and number of cells in each acinus was higher in the AMPK-inhibited (PRL+CC) condition compared to control (PRL+DMSO) acini (Figure 3J and 3K), consistent with the phenotype observed *in vivo* in AMPK KO mammary glands. Taken together, our findings demonstrate that AMPK is required for the maintenance of tissue architecture during mammary alveologenesis.

### AMPK loss enhances cellular proliferation during mammary alveologenesis

To assess whether the increased epithelial content in AMPK KO mammary glands is a result of hyperproliferation or reduced apoptosis, control and AMPK KO tissues were immunoprobed with the proliferation marker Ki67 and cyclin D1, and the apoptotic marker cleaved caspase-3. A significant increase in Ki67 staining (through immunohistochemistry) and elevated cyclin D1 levels (through immunoblotting) was observed in AMPK KO mammary glands at day 1 of lactation (Figure 4A and 4B). However, AMPK deletion did not affect proliferation at day 13.5 of pregnancy (Figure S4A). On the other hand, we did not observe any significant change in cleaved caspase-3 levels on AMPK ablation in day 1 lactating mammary glands as probed by immunoblotting and immunohistochemistry (Figure 4C and 4D). These data suggest that increase in epithelial content in the AMPK KO mammary gland is a consequence of hyperproliferation and not due to reduction in apoptosis.

**Figure 4.**
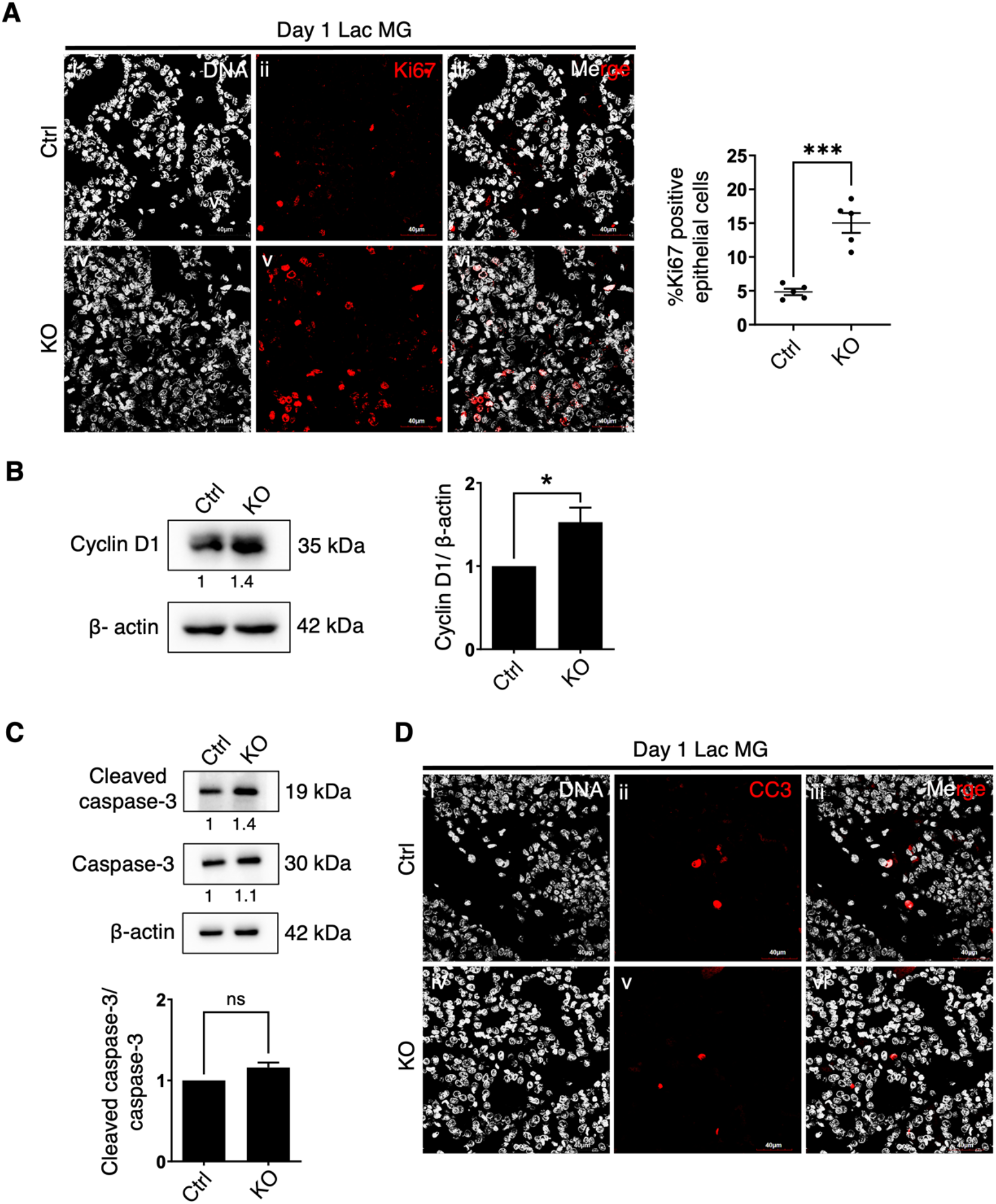
AMPK deletion and its effect on proliferation and apoptosis. (A) Laser confocal micrographs of control and AMPK KO mammary gland sections probed for Ki67 at day 1 of lactation show maximum intensity projection (MIP) of immunofluorescence signals from Ki67 (red) and staining of DNA with Hoechst (white); scale bar: 40 *μ*m. Representative scatter plots with bar graphs show the quantification of percentage Ki67 positive epithelial cells per field; error bars, mean ± SEM; *n* = 3. \*\*\**P*<0.001, unpaired Student’s *t*-test. (B and C) Representative immunoblots for specified proteins in control and AMPK KO mammary glands at day 1 of lactation. Graphs represent densitometric quantification of immunoblots; error bars, mean ± SEM; *n* = 3. \**P*<0.05, unpaired Student’s *t*-test, ns, not significant. (D) Laser confocal micrographs of control and AMPK KO mammary gland sections probed for cleaved caspase-3 (CC3) at day 1 of lactation show maximum intensity projection (MIP) of immunofluorescence signals from CC3 (red) and staining of DNA with Hoechst (white); scale bar: 40 *μ*m. (See also supplementary figure 4).

### AMPK ablation results in increased milk synthesis during pregnancy and lactation

Since we observed elevated AMPK expression and activity in wild-type lactating mammary glands (Figure 1), to determine whether the loss of AMPK affects lactating mammary gland function, we examined AMPK KO female mice for their nursing efficiency. AMPK KO female mice were fecund and gave birth to pups of similar litter sizes, comparable to their respective control females. Despite normal nursing characteristics displayed by AMPK KO dams, their pups were bigger in size than those nursed by control dams (Figure 5A). When measured for bodyweight, pups nourished by AMPK KO dams gained weight significantly faster from lactation day 7 onwards; the difference became exaggerated by day 14 lactation (Figure 5B and 5C). Furthermore, pups raised by AMPK KO dams showed a higher body weight than pups from control dam when examined at day 30 and day 60 postpartum (data not shown). To further investigate the reason for the increase in pups body weight, we first analysed the expression of major milk protein beta-casein at day 13.5 of pregnancy and day 1 of lactation. Immunoblot analysis showed a slight increase at day 13.5 of pregnancy which increased significantly at day 1 of lactation in AMPK KO mammary glands as well as in epithelial cells cultured as *ex vivo* monolayers (Figure S5A, 5D and S5B). Further, immunohistochemical analysis revealed an increase in beta-casein protein expression in KO mammary gland epithelia compared to control counterparts at day 1 of lactation (Figure 5E), suggesting an inhibitory role of AMPK in milk protein synthesis. To further confirm the effect of AMPK inhibition on beta-casein protein expression, we cultured wild-type primary MECs on LrECM substrata for four days in the presence of CC. In the presence of prolactin, CC treatment increased beta-casein synthesis, indicating that increased beta-casein synthesis previously observed *in vivo* is likely a direct consequence of AMPK deletion in luminal cells (Figure 5F, S5C, and 5G).

**Figure 5.**
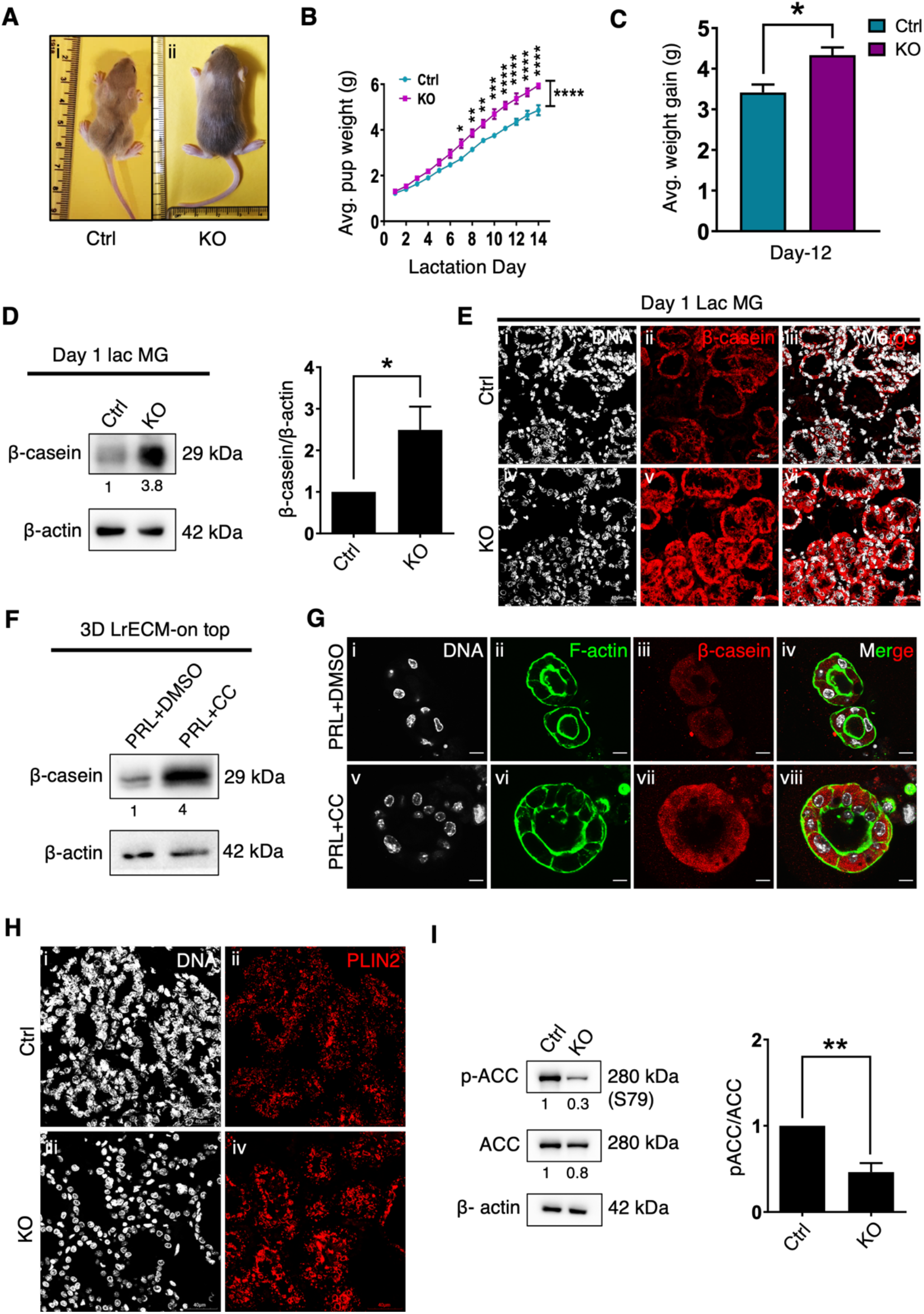
Effect of AMPK deletion on milk protein and lipid expression. (A) Representative picture of bigger size pup nursed by AMPK KO dam (right) compared to pup nursed by control dam (left) on postnatal day 14. (B) Growth-rate analysis of pups nursed by control and AMPK KO lactating mice; litter size is 5 per set; error bars, mean ± SEM; *n* = 3. \**P*<0.05, \*\**P*<0.01, \*\*\**P*<0.001, \*\*\*\**P*<0.0001, unpaired Student’s *t*-test and two-way ANOVA test. (C) Bar graph shows average weight gain by pups after 12 days of birth; error bars, mean ± SEM; *n* = 3. \**P*<0.05, unpaired Student’s *t*-test. (D) Representative immunoblots for specified proteins in control and AMPK KO mammary glands at day 1 of lactation. Graph represents densitometric quantification of immunoblots; error bars, mean ± SEM; *n* = 4. \**P*<0.05, unpaired Student’s *t*-test. (E) Laser confocal micrographs of control and AMPK KO mammary gland sections probed for β-casein at day 1 of lactation show maximum intensity projection (MIP) of immunofluorescence signals from β-casein (red) and staining of DNA with Hoechst (white); scale bar: 40 μm. (F) Representative immunoblots for specified proteins in MECs cultured on-top of LrECM for 5 days in the presence of PRL treated with vehicle control DMSO and CC. (G) Laser confocal z-stack micrographs of MECs cultured on-top of LrECM for 5 days in the presence of PRL treated with vehicle control DMSO (i-iv) and compound C (CC) (v-viii), show maximum intensity projection (MIP) of immunofluorescence signals from β-casein (red), staining of F-actin with phalloidin (green), and DNA with Hoechst (white); scale bar: 10 *μ*m. (H) Laser confocal micrographs of control and AMPK KO mammary gland sections probed for PLIN2 at day 1 of lactation show maximum intensity projection (MIP) of immunofluorescence signals from PLIN2 (red) and staining of DNA with Hoechst (white); scale bar: 40 *μ*m. (I) Representative immunoblots for specified proteins in control and AMPK KO mammary glands at day 1 of lactation. Graphs represent densitometric quantification of immunoblots; error bars, mean ± SEM; *n* = 3. \*\**P*<0.01, unpaired Student’s *t*-test. (See also supplementary figure 5).

To assess the role of AMPK in lipogenesis and secretory activation, we analysed the status of lipid droplet-associated protein perilipin 2 (PLIN2). Immunostaining for PLIN2 in AMPK KO mammary glands showed a greater number of lipid droplets in luminal epithelia at day 1 of lactation (Figure 5H), suggesting enhanced lipid synthesis on AMPK deletion. Phosphorylation of ACC at Ser79 by AMPK inhibits fatty acid synthesis in bovine and goat MECs (McFadden and Corl, 2009; Zhang et al., 2011). Herein, we found reduced ACC phosphorylation at Ser79 in AMPK KO mammary glands, corroborating increased lipid synthesis on AMPK ablation (Figure 5I and 2F). Taken together, these data suggest that AMPK regulates milk synthesis during lactation in mammary gland and knocking out AMPK leads to increased milk synthesis.

### AMPK ablation leads to enhanced STAT5 phosphorylation

Prolactin regulates mammary alveolar morphogenesis and differentiation via PRL-JAK2-STAT5 signaling pathway (Ormandy et al., 1997). PRL binding to prolactin receptor (PRLR) activates cytoplasmic STAT5 via JAK2-mediated phosphorylation at tyrosine residue (Y694) (Gouilleux et al., 1994; Liu et al., 1995). Overactivation of STAT5 in murine mammary gland leads to increased cell proliferation, dense mammary alveolar structures, and overexpression of beta-casein (Iavnilovitch et al., 2002). Our observations indicated that AMPK knockout phenocopies STAT5 activation; therefore, we investigated whether the PRL-JAK2-STAT5 signaling is perturbed upon AMPK deletion. To assess this, we first analysed STAT5 expression and activity (pSTAT5^Y694^) in control and AMPK KO mammary glands. Immunoblot analysis of mammary tissue extracts at day 13.5 of pregnancy did not show any change in STAT5 and pSTAT5^Y694^ levels (Figure 6A). Immunoblot analysis of AMPK KO mammary glands at day 1 of lactation revealed unchanged total STAT5 but elevated pSTAT5^Y694^ levels when compared with control tissues (Figure 6B). However, signals for PRLR and pJAK2^Y1007/1008^ were unchanged in AMPK KO lactating gland tissues (Figure 6C-6F), indicating the increase in STAT5 phosphorylation in the absence of AMPK is not due to altered PRLR expression and JAK2 phosphorylation. Taken together, our findings suggest that AMPK indirectly modulates STAT5 activation and, in turn, its target genes expression to bring about regulated alveolar development during pregnancy and lactation.

**Figure 6.**
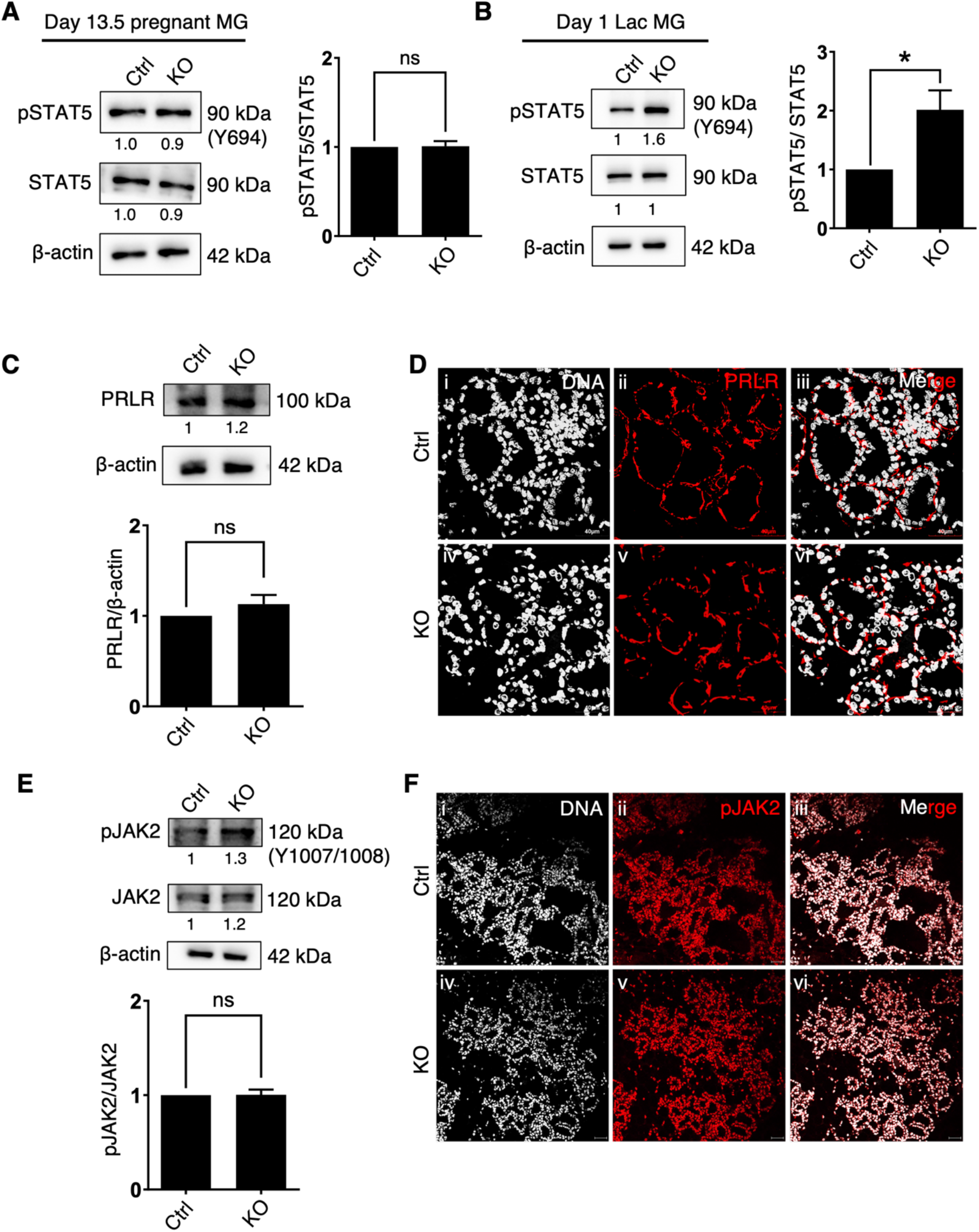
AMPK regulates STAT5 phosphorylation in mammary gland. (A-C) Representative immunoblots for specified proteins in control and AMPK KO mammary glands at day 13.5 of pregnancy (A) and day 1 of lactation (B and C). Graphs represent densitometric quantification of immunoblots; error bars, mean ± SEM; *n* = 5. \**P*<0.05, unpaired Student’s *t*-test, ns, not significant. (D) Laser confocal micrographs of control and AMPK KO mammary gland sections probed for PRLR at day 1 of lactation show maximum intensity projection (MIP) of immunofluorescence signals from PRLR (red) and staining of DNA with Hoechst (white); scale bar: 40 *μ*m. (E) Representative immunoblots for specified proteins in control and AMPK KO mammary glands at day 1 of lactation. Graph represents densitometric quantification of immunoblots; error bars, mean ± SEM; *n* = 3. ns, not significant. (F) Laser confocal micrographs of control and AMPK KO mammary gland sections probed for pJAK2 at day 1 of lactation show maximum intensity projection (MIP) of immunofluorescence signals from pJAK2 (red) and staining of DNA with Hoechst (white); scale bar: 40 *μ*m.

## Discussion

This study reveals a novel role of AMPK in mammary epithelial growth and differentiation during pregnancy and lactation. High AMPK expression and activity was found in pregnant and lactating murine mammary glands and in MECs cultured on LrECM scaffolds in the presence of prolactin. Genetic ablation of AMPK in mammary glands led to an increase in STAT5 phosphorylation during lactation. AMPK KO MECs were hyperproliferative and showed altered lactogenic differentiation during alveolar morphogenesis. In addition, there was increased milk production and improved nursing efficiency of lactating dams on AMPK ablation. These data begin to integrate crosstalk among hormonal cues, metabolic regulators, and signaling pathways in co-ordinating mammary gland alveologenesis and lactation.

The physiological purpose of the mammary gland is to synthesize and secrete milk during lactation to provide nourishment to the neonates. The synthesis of major milk components such as protein, lipid, and carbohydrate in MECs requires an adequate energy supply (Huang et al., 2020). In order to meet the energy demands for milk synthesis and tissue homeostasis, various catabolic pathways get activated to balance the ATP production in response to higher ATP utilisation (Burgos et al., 2013). AMPK activation has been reported in bovine MECs upon energy deprivation in monolayer cultures (Zhang et al., 2018). Recent data suggest a major role for prolactin in energy and metabolic homeostasis (Perez Millan et al., 2014; Tovar and Diéguez, 2014). Our study builds on these prior observations and provides insights into hormonal induction of AMPK during lactation. In contrast to monolayer cultures, we found a significant increase in AMPK expression and activity when murine MECs were grown on LrECM scaffolds and allowed to form lumen-containing acini in the presence of PRL. This contextual increase in AMPK expression in response to prolactin can be attributed to the basally located PRL receptors, which are otherwise inaccessible to ligands in monolayers but are rendered accessible when multicellular polarity emerges in 3D (Xu et al., 2009), suggesting that increased AMPK expression and activity is driven by PRL in a LrECM permissive manner. Our study, for the first time, links AMPK signaling with the well-studied prolactin-JAK2/STAT5 pathway in the regulation of acinar morphogenesis. However, the molecular crosstalk between these pathways is still elusive and will be addressed in future studies.

Our observations reveal a novel homeostatic role for AMPK in cellular proliferation with implications in breast cancer. We found increased epithelial cell proliferation with a significant increase in cytokeratin 14 and 18 expressing luminal and myoepithelial cells, respectively, in AMPK KO mammary glands. These data correlate with earlier findings of mammary gland hyperplasia observed in Cyclin D1 overexpressing transgenic mice (Wang et al., 1994). In addition, aberrant activation of STAT5 has been shown to be weakly oncogenic in mouse model of breast cancer (Iavnilovitch et al., 2002). In the absence of AMPK, we found an increase in the tyrosine-694 phosphorylation and activity of STAT5 with a corresponding increase in beta-casein protein expression. These data suggest that, in a normal scenario, AMPK may keep STAT5 phosphorylation in check and thereby regulates PRL-JAK2-STAT5 signaling. However, the mechanism through which AMPK affects STAT5 is still unclear and needs further investigation. It is interesting to note that while prolactin induces AMPK activity during acinar morphogenesis, loss of AMPK leads to increased epithelial proliferation and altered lactogenic differentiation *in vivo*. Our findings suggest that prolactin signaling may therefore self-regulate itself in an AMPK-dependent manner. This is reinforced by our findings that AMPK signaling decreases STAT5 activation. Collectively, our data suggest that AMPK activity is required to bring about regulated lobuloalveolar expansion and functional differentiation. In the absence of AMPK, there is an aberrant lobuloalveolar expansion which may drive the gland towards tumor formation, thereby supporting the role of AMPK as a metabolic tumor suppressor. However, once tumor is initiated, we and others have shown tumor-promoting roles of AMPK by enhancing breast cancer cell survival under various stress conditions (Hindupur et al., 2014; Jeon and Hay, 2015; Ng et al., 2012; Saha et al., 2018; Sundararaman et al., 2016), as well as by promoting EMT (Saxena et al., 2018) and stemness (Lahiry et al., 2020). Recent murine models have further elaborated on this dual role of AMPK in cancer (Vara-Ciruelos et al., 2020).

Milk is mainly composed of three proximal nutritional constituents: proteins, lipids, and lactose. Synthesis of these components during lactation requires abundant energy (Dewey, 1997). In our study, effect of AMPK ablation was mainly observed during the differentiation phase of pregnant and lactating mammary glands. We found a slight increase in beta-casein levels as early as day 13.5 of pregnancy, which increased significantly during lactation in AMPK KO glands. This was further confirmed by analysing the expression of beta-casein in 3D cultured MECs upon AMPK inhibition. Fatty acids and other lipid components are major constituents required for milk lipid synthesis. ACC is the principal catalytic enzyme involved in fatty acid synthesis and regulates lipogenesis in animal tissues (Gibson et al., 1958). AMPK activation induces ACC phosphorylation, thereby inhibiting fatty acid synthesis in bovine and goat MECs, hence, conserving ATP (McFadden and Corl, 2009; Zhang et al., 2011). In our study, we found a significant decrease in ACC phosphorylation in the AMPK KO mammary glands, suggesting an increase in lipid synthesis. AMPK-mediated phosphorylation of milk lipid surface protein, perilipin (PLIN), triggers lipolysis (Kaushik and Cuervo, 2016). We found increased PLIN2 expressing lipid droplets in AMPK-null mammary tissues. These findings suggest that AMPK regulates milk protein and lipid synthesis during lactation, further affecting pups weight. AMPK is shown to suppress global protein synthesis by inhibiting mTOR in bovine MECs (Burgos et al., 2013). In line with the earlier findings, we suggest that increased milk protein expression upon AMPK deletion could be via regulation of mTOR pathway. Since we found an increase in both AMPK activation and milk protein synthesis in lactating mammary gland, it would be interesting to understand the translational machinery of milk protein synthesis and associated role of AMPK-mTOR signaling pathway in its regulation.

In summary, our findings show that AMPK plays a homeostatic role during pregnancy and lactation by regulating epithelial cell proliferation and lactogenesis. In addition, this study places AMPK at a central node upstream of STAT5 activation providing evidence for its diverse control of various cellular processes during lactation.

## Materials and methods

### Mouse model

Floxed *PRKAA1* (AMPKα1) and floxed *PRKAA2* (AMPKα2) animals have been generated as described previously (Boudaba et al., 2018; Viollet et al., 2003). Animals having both floxed *PRKAA1* (AMPKα1) and floxed *PRKAA2* (AMPKα2) were generous gifts from Drs. Marc Foretz and Benoit Viollet, Université de Paris. The MMTV-Cre (line D) mice were purchased from the Jackson Laboratory (Stock No: 003553). All animals were maintained on the mixed background of FVB/N and C57Bl/6J. For all studies, littermate control (AMPKα1^fl/fl^, AMPKα2^fl/fl^) and conditional knockout (AMPKα1^-/-^, AMPKα2^-/-^) mice were used. In the AMPK floxed mice, LoxP sites were inserted in introns flanking exon 4 and exon 5 of *PRKAA1* encoding amino acids 113-190 and exon 6 of *PRKAA2* encoding amino acids 189–260. In the presence of Cre recombinase, nucleotide sequences encoding parts of the catalytic domain of *PRKAA1* and *PRKAA2* are removed. The presence of floxed *PRKAA1* and floxed *PRKAA2* and Cre allele was confirmed by PCR genotyping of DNA samples isolated from tail biopsy. Primers used for PCR are listed in Table 1. All mice were mated at around 8-10 weeks of age. Pups growth analysis was performed by maintaining litter sizes of 5 pups/litter. Pups were weighed every day till 14 days of lactation and at postnatal day 30^th^ and 60^th^. All experiments involving animals were reviewed and approved by the Institutional Animal Ethics Committee of Indian Institute of Science, Bangalore. The mice were maintained on a 12 hours light and 12 hours dark cycle in the animal facility and fed with standard rodent chow diet (Altromin) and RO purified water.

### Antibodies and pharmacological reagents

The following primary antibodies were used in the study: AMPKα (CST, #2532), AMPKα1 (CST, #2795), AMPKα2 (CST, #2757), pAMPKα^T172^ (CST, #2535), ACC (CST, #3662), pACC^Ser79^ (CST, #3661), β-Actin (Invitrogen, #MA-140), Cre recombinase (CST, #15036), Cyclin D1 (ABclonal, #A19038), Ki67 (Abcam, #ab15580), Cleaved caspase-3 (CST, #9664), Caspase-3 (CST, #9662), STAT5 (ABclonal, #A7733), pSTAT5^Tyr694^ (CST, #9359 and Invitrogen #71-6900), Beta-casein (Santa Cruz, #SC-166530 and #SC-30041), CK-14 (Abcam, #ab7800), CK18 (Abcam, #ab668), JAK2 (ABclonal, #A11497), pJAK2^Y1007/1008^ (Abcam, #ab219728), PRLR (Abcam, #ab170935 and Invitrogen, #35-9200), PLIN2 (ABclonal, #A6276). Secondary antibodies used for the study were anti-mouse IgG conjugated with cy3 (Abcam, #ab97035), anti-mouse IgG Alexa Fluor 647 (Invitrogen, #A21236), antirabbit IgG conjugated with Alexa Fluor 568 (Invitrogen, #A11011), anti-rabbit IgG Alexa Fluor 647 (Invitrogen, #A27040). Nucleus was counterstained by Hoechst 33342 (Sigma, #B-2261), and F-actin was stained by Phalloidin Alexa Fluor 488 (Invitrogen, #A12379). Pharmacological inhibitor of AMPK, compound C (Dorsomorphin), Calbiochem #171260, was used in this study at 5 μM concentration. DMSO (Dimethyl sulfoxide), Calbiochem #317275, was used as vehicle control.

### Isolation of mammary gland, whole mount analysis, and H&E staining

The 4^th^ inguinal mammary glands of female mice were dissected from different stages of mammary gland development and spread on positively charged adhesive slides (Hisure scientific, #9023). Whole mounts were fixed overnight in Carnoy’s fixative solution (60% ethanol, 30% chloroform, and 10% glacial acetic acid). For whole mount analysis, slides were stained with carmine-alum overnight, dehydrated by passing slides in ethanol series (50%, 70%, 90%, and 100%: 2 hours for each step). Slides were cleared in methyl salicylate and mounted using DPX mountant (Fisher Scientific, #18404). Images of whole mounts were captured at different magnifications using stereoscope (Magnus MSZ-TR).

For H&E staining and sections preparation, Carnoy’s solution fixed mammary glands were paraffin-embedded, and 5 μm thin sections were prepared. These sections were transferred on adhesive charged glass slides and stained for hematoxylin (HiMedia #S034) and eosin (SDFCL #45380) for histological analysis, and the rest unstained sections were used for IHC analysis.

### Immunohistochemistry and Immunofluorescence

5 μm thin tissue sections were deparaffinized in oven at 60°C for 2 hours to overnight and washed in xylene for 3 changes of 15 minutes each step. After deparaffinization, sections were rehydrated by passing through gradient of ethanol. For antigen retrieval, sections were boiled in 0.01 M Tris-EDTA buffer (pH – 9.0) for 15 minutes. After cooling sections, endogenous peroxidase activity was blocked by incubation of slides in 3% H2O2 for 30 minutes. After blocking for 45 minutes, sections were incubated with primary antibodies at 4°C overnight. Then slides were incubated in enhancer solution and polymer HRP conjugated secondary antibody as instructed by the manufacturer (BioGenex, Supersensitive^™^ IHC detection kit, #QD430-XAKE). DAB chromogen was added to each section for 10 minutes, and slides were washed three times with TBS buffer containing 0.05% Tween-20. Sections were counterstained with hematoxylin for 1 minute and mounted using DPX mountant. Images were captured at different magnifications in bright-field microscope (Olympus IX-71).

For immunofluorescence analysis, sections were permeabilized with 0.5% Triton-X 100 for 10 minutes after antigen retrieval. Sections were incubated with primary antibody overnight at 4°C, and slides were washed three times in PBS buffer containing 0.05% Tween-20 and incubated with fluorochrome-conjugated secondary antibodies for 2 hours at room temperature. Sections were counterstained with Hoechst dye for 5-10 minutes and mounted with mounting media containing anti-fading reagent. Slides were then imaged in Confocal microscope (Olympus FluoView FV10i and Leica TCS SP-8). Images were processed using ImageJ software (NIH).

### Immunocytochemistry

Monolayered and 3D cultured acini were fixed in 3.7% formaldehyde at room temperature for 10 minutes and 20 minutes, respectively. Cells were permeabilized with 0.5% Triton-X 100 in PBS for 10 minutes and blocked for 1 hour in 3% BSA. Primary antibodies were incubated in blocking buffer overnight at 4°C. Cells were washed with PBST and stained with secondary antibody (1:500) following day. Cells were counterstained with Hoechst (1:1000) and mounted. Images were captured using epifluorescence microscope (Olympus IX71) and confocal microscope. Images were processed and analysed using ImageJ software.

### Immunoblotting

Isolated mammary glands were homogenized in 1 x RIPA lysis buffer containing 150 mM sodium chloride, 1% NP40, 0.5% sodium deoxycholate, 0.1% SDS, 50 mM Tris, supplemented with 1 x protease inhibitor and 1 x phosphatase inhibitor (1 mM PMSF, 5 mM EDTA, 10 mM NaF, and 1 mM Na_2_OV_4_) and incubated on ice for 20 minutes. Protein supernatant was obtained by precipitating cell debris by centrifugation in a bench-top centrifuge at 13000 rpm for 10 minutes at 4^°^C. For Matrigel embedded 3D cell culture, cells were first scrape-lysed in chilled PBS-EDTA buffer and then lysed using RIPA lysis buffer, homogenized, and centrifuged to remove Matrigel as described earlier (Mroue and Bissell, 2013). Protein concentrations were determined by Bradford assay and Nanodrop method. An equal amount of protein (50-100 μg in each well) was loaded and resolved by SDS-polyacrylamide gel electrophoresis. After resolving, proteins were transferred to PVDF membrane. For proteins probing, PVDF membrane was blocked in 5% BSA for 45 minutes at room temperature followed by incubation with primary antibody overnight at 4^°^C. After 3 TBST washes, membrane was incubated with secondary antibody conjugated with HRP (Jackson Laboratory) at room temperature for 2 hours. Blots were developed by ECL substrate (1:1; H_2_O_2_: Luminol) and visualized under Gel Doc system (Syngene-G box). Multipanel blots were compiled by either reprobing the same blot for successive antibodies, or by running the same lysate multiple times from a master-mix of prepared sample; β-actin served as loading control for each run. Representative immunoblots show data consistent with minimally three independent experiments. Blot intensity was measured by densitometric analysis in ImageJ software.

### Primary mammary cell isolation and 3D culture

The 4^th^ inguinal mammary gland was dissected from control and AMPK KO female mice at 13-16 days of pregnancy. Tissues were minced using a surgical blade and homogenized. Minced tissues were digested at 37°C by gently shaking with DMEM/F12 (Sigma, #D-8900) media supplemented with 0.2% collagenase (Sigma, #C-2674), 0.2% trypsin (Sigma, #T-4799), 5% fetal bovine serum (Gibco, #10270-106) and 50 *μ*g/ml penicillin (HiMedia, #TC-020), and streptomycin (HiMedia, #TC-305). Epithelial organoids were pelleted by centrifugation at 1500 rpm for 10 minutes and re-suspended in 4 ml of DMEM/F12 with 40 *μ*l of DNase (Sigma, #5025) (2 U/*μ*l). Isolated organoids were cultured as monolayer in primary mammary epithelial culture media [DMEM/F12 1:1, supplemented with 50 *μ*g/ml hydrocortisone (Sigma, #H0888), 10 mg/ml insulin (Sigma, #I9278), and 5 *μ*g/ml EGF (Sigma, #E9644)] and trypsinised for organotypic culture. 1 × 10^4^ cells were seeded on-top of solidified Laminin-rich extra cellular matrix (LrECM) (commercially available as Matrigel, Corning, #354230) in 8-well chamber/ 96-well plate for immunocytochemical analysis, and 1 × 10^5^ cells/cm^2^ were seeded for protein lysate preparation. After 24 hours of seeding, cells were allowed to differentiate for 5 days with differentiating media [3 μg/ml prolactin (Sigma, #L6520, and Dr. A.F. Parlow, Lundquist Institute/ NHPP), 10 mg/ml insulin, 50 μg/ml hydrocortisone] with and without compound C at 5 μM concentration, replenishing media on alternative day.

### Statistical analysis

Data are represented as mean ±S.E.M. P values were determined by Student’s unpaired t-test or two-way ANOVA using GraphPad Prism 8.0.2 software. The significance level is indicated by different numbers of asterisks as follows: \**P*<0.05, \*\**P*<0.01, \*\*\**P*<0.001, and \*\*\*\**P*<0.0001.

## Acknowledgement

This work was supported by grants from the Indian Institute of Science (IISc), Bangalore, India, the Department of Biotechnology (Govt. of India)-IISc partnership programme, the International Development Research Centre, IDRC, and the Indo-Israel Joint Scientific Research Program funded by University Grants Commission (UGC), Govt. of India. Support from DST-FIST and UGC, Government of India, to the Department of MRDG are also acknowledged. All student fellowships and faculty salaries are through MHRD funding through IISc. The studies described here were carried out according to the guidelines of the institutional review board and Institutional Animal Ethics Committee (IAEC) (CAF/ Ethics/ 845/2021). Authors acknowledge the Central Animal Facility (CAF), IISc, for animal work and the Biological Sciences Imaging facility for microscopy.

## Author Contribution

S.J. conceived and conceptualized the study, performed most experiments, undertook data analyses, compiled figures, and wrote the manuscript. P.S. and K.C. helped with experimentation, data acquisition, analyses, and manuscript writing. S.M. contributed experiments. MF and BV developed and gifted AMPK floxed mice, and edited the manuscript. BV and NR supervised generation of conditional AMPK KO mice and edited manuscript. R.B. contributed ideas, supervised, and edited the manuscript. A.R. contributed ideas, project administration, funding acquisition, supervision, and edited the manuscript.

---

**Supplementary Figure 1.**
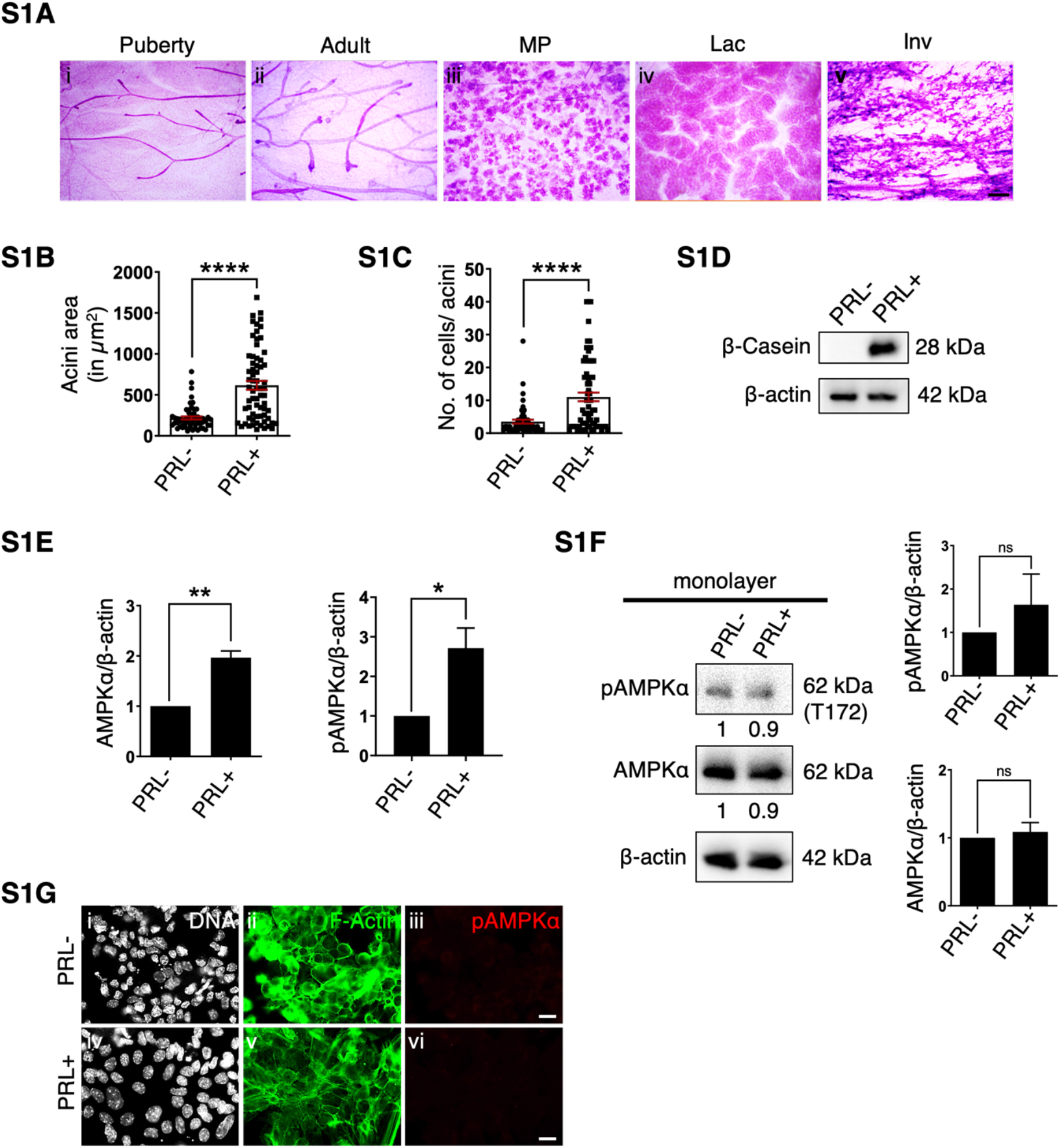
Expression and activity of AMPK across different stages of mammary gland development. (A) Micrographs of carmine-stained whole mounts of mammary glands at puberty (5 weeks), adult (10 weeks), MP (mid-pregnant), Lac (day 7 lactation), and Inv (day 7 involution). (B-D) Representative scatter plots with bar graphs (B and C) show the quantification of area and total number of cells per acinus; error bars, mean ± SEM; *n* = 3. \*\*\*\**P*<0.0001, unpaired Student’s *t*-test and immunoblots (D) for specified proteins in primary MECs cultured on-top of LrECM scaffolds in the presence (+) and absence (-) of PRL for 5 days. (E) Graphs represent densitometric quantification of immunoblots (Figure 1E); error bars, mean ± SEM; *n* = 4. \**P*<0.05, \*\**P*<0.01, unpaired Student’s *t*-test. (F and G) Representative immunoblots (F) and laser confocal micrographs of immunostaining (G) for specified proteins in primary MECs cultured as monolayer with and without PRL. Graphs represent densitometric quantification of immunoblots; error bars, mean ± SEM; *n* = 2. ns, not significant. Micrographs show maximum intensity projection (MIP) of immunofluorescence signals from pAMPKα^T172^ (red), staining of F-actin with phalloidin (green), and DNA with Hoechst (white); scale bar: 20 μm.

**Supplementary Figure 2.**
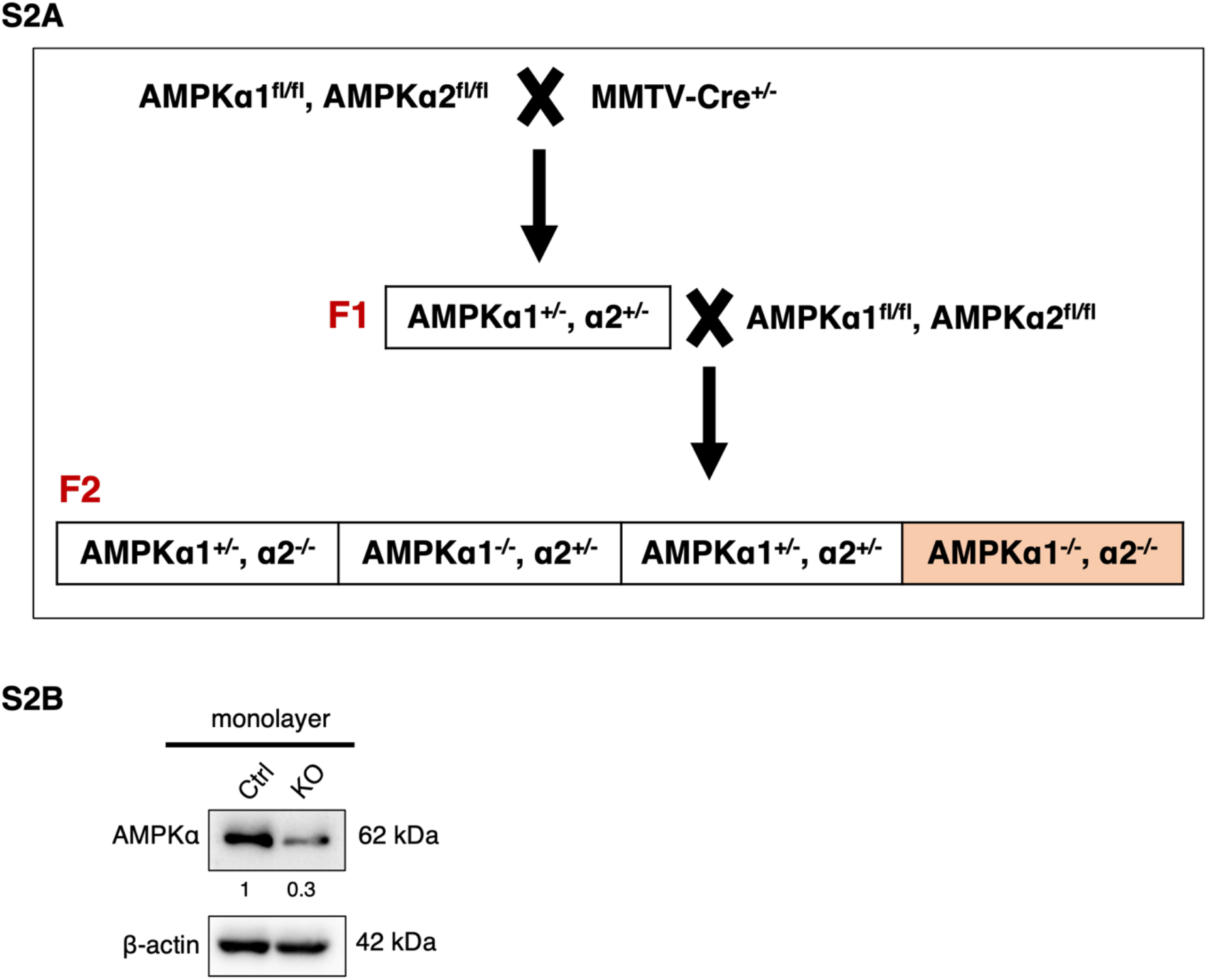
Generation of AMPKα conditional knockout mice. (A) Schematic representation of generation of AMPK heterozygous KO mice in F1 generation and of AMPK homozygous KO mice in F2 generation. (B) Representative immunoblots for specified proteins in control and AMPK KO primary MECs cultured as monolayer.

**Supplementary Figure 3.**
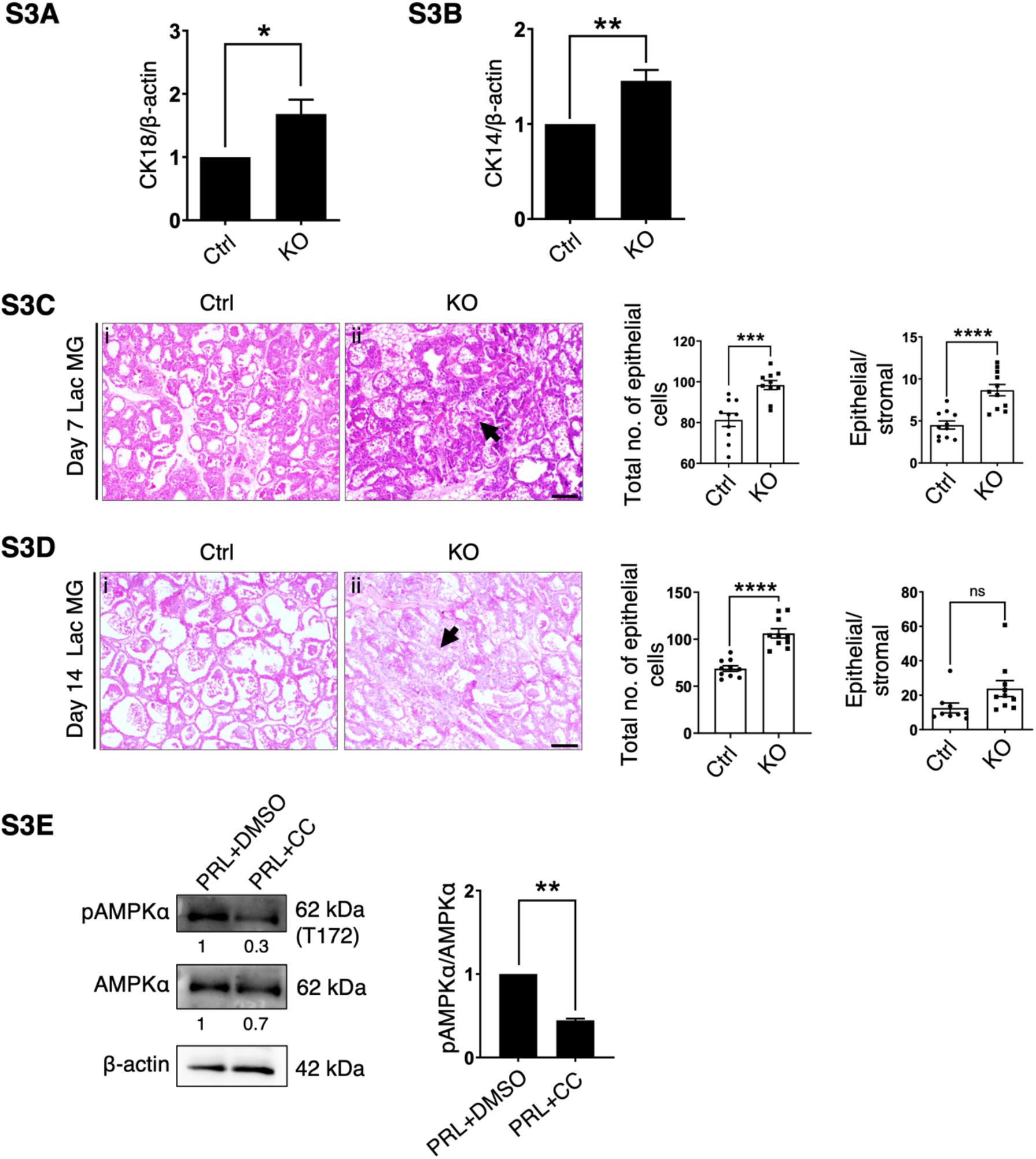
Effect of AMPK ablation on lobuloalveolar morphogenesis. (A and B) Graphs represent densitometric quantification of immunoblots (Figure 3G and 3H); error bars, mean ± SEM; *n* = 3. \**P*<0.05, \*\**P*<0.01, unpaired Student’s *t*-test. (C and D) Micrographs of hematoxylin and eosin (H&E) stained sections of control and AMPK KO mammary glands at day 7 (C) and day 14 (D) of lactation. Arrow indicates increased epithelial content in AMPK KO mammary glands; scale bar: 50 *μ*m. Representative scatter plots with bar graphs show the quantification of total number of epithelial cells per field and epithelial/ stromal content in control and AMPK KO mammary glands at day 7 (C) and day 14 (D) of lactation (at least 10 random fields per gland); error bars, mean ± SEM; *n* = 3. \*\*\**P*<0.001, \*\*\*\**P*<0.0001, unpaired Student’s *t*-test, ns, not significant. (E) Representative immunoblots for specified proteins in MECs cultured in LrECM for 5 days in the presence of PRL treated with vehicle control DMSO and CC. Graphs represent densitometric quantification of immunoblots; error bars, mean ± SEM; *n* = 3. \**P*<0.05, unpaired Student’s *t*-test.

**Supplementary Figure 4.**
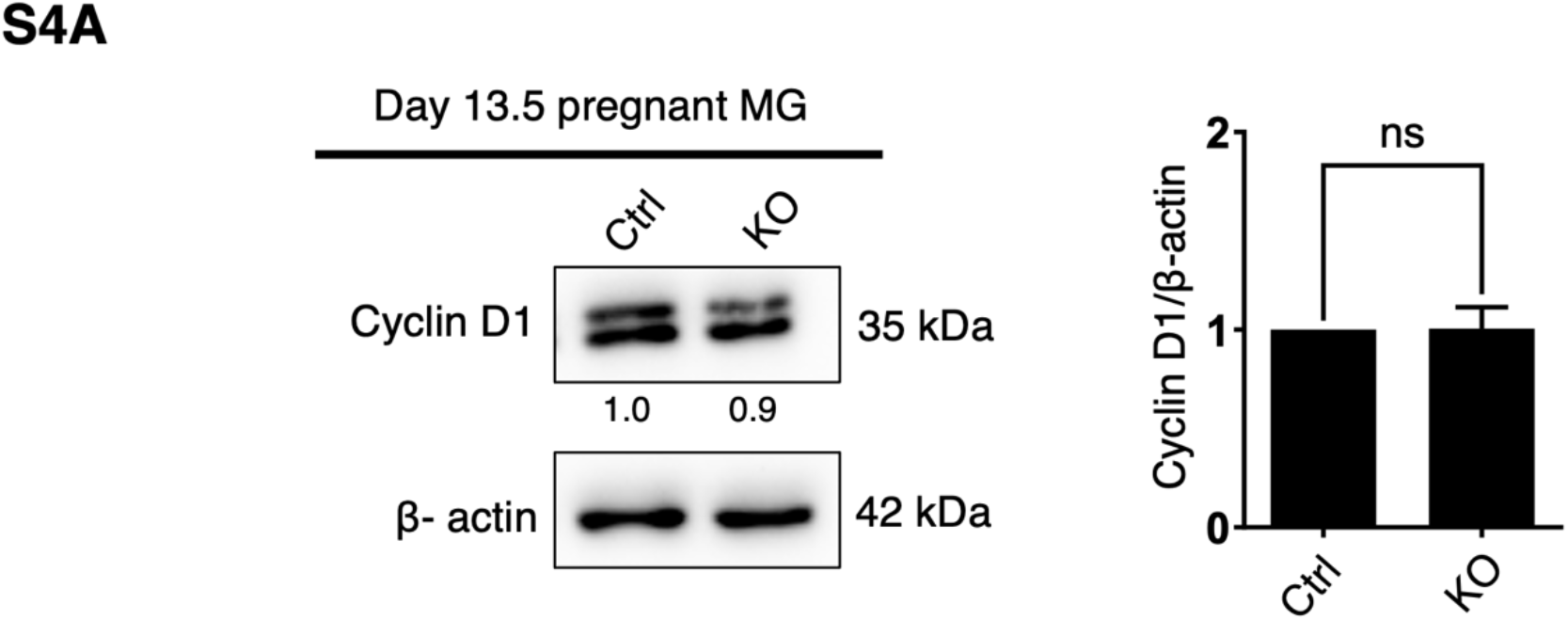
AMPK deletion and its effect on proliferation and apoptosis. (A) Representative immunoblots for specified proteins in control and AMPK KO mammary glands at day 13.5 of pregnancy. Graphs represent densitometric quantification of immunoblots; error bars, mean ± SEM; *n* = 3. ns, not significant.

**Supplementary Figure 5.**
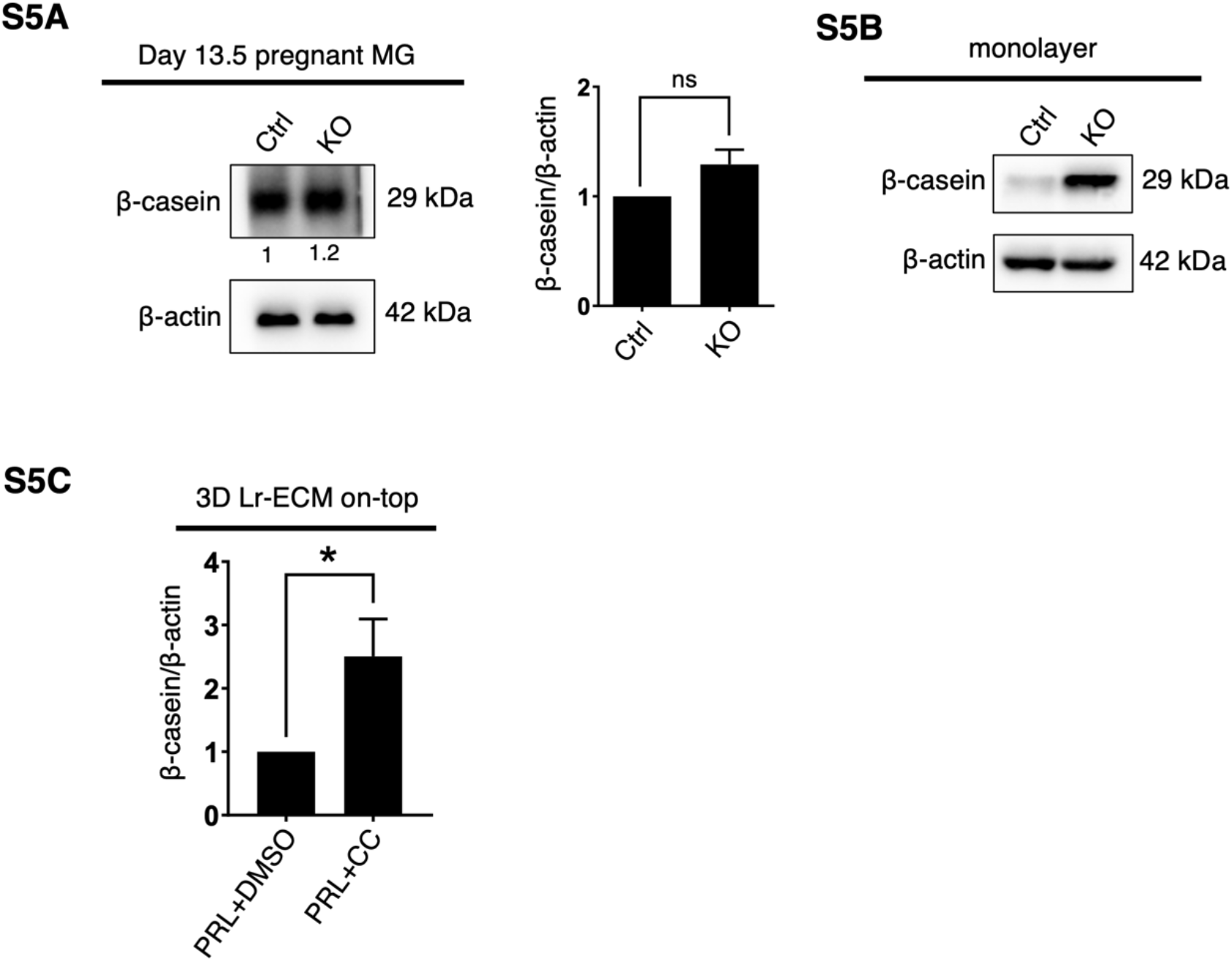
Effect of AMPK deletion on milk protein and lipid expression. (A-C) Representative immunoblots for specified proteins in control and AMPK KO mammary glands at day 13.5 of pregnancy (A), in control and KO MECs cultured as monolayer (B), and in MECs cultured in LrECM for 5 days in the presence of PRL treated with vehicle control DMSO and compound C (CC) (C). Graphs represent densitometric quantification of immunoblots; error bars, mean ± SEM; *n* = 3 (A), *n* = 4 (C). \**P*<0.05, unpaired Student’s *t*-test, ns, not significant.

